# A novel deep optimal transport framework reveals prostate cancer risk heterogeneity, Alzheimer’s disease risk heterogeneity, and myeloma cells associated with a short-term sham bortezomib response and progressive disease

**DOI:** 10.1101/2025.02.03.636036

**Authors:** Ziyu Liu, Sihong Li, Justin Couetil, Jie Zhang, Kun Huang, Travis S. Johnson

**Affiliations:** Purdue University; Indiana University School of Medicine

**Keywords:** Single cell RNA sequencing, Spatial Transcriptomics, Alzheimer’s disease, Prostate cancer, Multiple myeloma, Transfer learning, Deep learning, Multitask learning, Wasserstein distance, Optimal Transport

## Abstract

Leveraging single-cell gene expression profiles can significantly enhance our understanding of diseases by associating single cells with traits such as disease subtypes, prognosis, and drug response. Although previous efforts have linked single cell clusters and groups with these attributes, they have primarily focused on changes in cell proportions while overlooking transcriptional changes at the single cell level. To further unravel cell heterogeneity with clusters and reveal the nuanced behaviors of cellular subtypes, it is essential to assess the disease associations of individual cells. Previous methods often fail to capture complex patterns that are only discernible through summarizing non-linear relationships across multiple genes. The Diagnostic Evidence GAuge of Single-cells/Spatial-transcriptomics (DEGAS) framework advances these efforts by aligning single cells and/or spatial transcriptomics regions with patients through a unified latent space using non-linear transformations learned from deep neural networks (DNNs). DEGAS achieves superior performance in analyzing single cell and spatial transcriptomics datasets, including Alzheimer’s disease (AD), multiple myeloma (MM), and prostate cancer (PDAC). Here we present DEGAS version 2, which has been updated with optimial transport based transfer learning and improved time-to-event loss functions, more advanced model architecture, and improved model baseline evaluations. DEGASv2 outperformed other methods in both single cell and spatial transcriptomic baseline comparisons. On the multiple myeloma discovery dataset, DEGASv2 enabled us to discover cell types that exhibited distinct drug response patterns over various time frames and were validated with time series single cell multiomic data that we generated, demonstrating a dangerous subtype of cell and novel therapeutic target.

## INTRODUCTION

Single cell (SC) sequencing techniques allows us to measure gene expression on a cellular level and has vastly broadens our comprehension of cellular profiles in various human organs and tissues [1], This advancement has improved our insight into a wide array of diseases, ranging from cancers to diabetes [2]. Significant strides have been made to associate SC clusters/groups with disease-relevant traits, such as subtypes, prognosis, and drug responsiveness [3]. These intensive efforts uncover the relationship between cells and disease based on variations in cell proportion/abundance or transcriptional changes. For instance, [4, 5] applied SC expression deconvolution techniques to observe shifts in cell subtype proportions across different disease states, thereby linking cell types to pertinent disease characteristics.

However, these methods have not fully harnessed transcriptional variations among cells. Consequently, several studies [6, 7] sought to connect SC clusters with patient-level attributes through differentially expressed genes (DEGs) associated with disease traits by assessing their prevalence in distinct SC groups. Nevertheless, the aforementioned methods do not offer comprehensive insights into all cellular subgroups because they correlate disease attributes with cell type- s/clusters as collective entities, thereby neglecting the heterogeneity within each cell type/cluster. As a result, the function and signal of individual cells may be obscured by the characteristics of the cluster to which they belong. To uncover subgroups of cells that behave differently from others within the same group/cluster, we aim to develop a tool that enables the investigation of the disease association score of each individual single cell.

Several attempts have been made to associate each individual SC with disease attributes by leveraging the similarity of gene expression profiles between SC and bulk patient samples [8–10]. This approach is rooted in the observation that bulk- level measurements, which aggregate RNA expression from all cells within a tissue, are directly correlated with patient-level disease traits in extensive datasets, such as the TCGA cohort [11]. Given that these cohorts typically contain data from hundreds of patients—a number considerably larger than that found in SC cohorts, where SCs are sampled from only a handful of patients—these bulk gene expression profiles are more amenable to statistical analyses, such as classification or regression. Building on this concept, the Scissor method [9] determined the association strength of each cell with the patient-level phenotype by fitting a linear regression model using the patient-to-cell Pearson correlation matrix as the design matrix and sample phenotype as the response variable.

Coefficients of the linear model was then used as the indicator of whether a cell was positively or negative associated with a trait. Yet, Scissor was unable to detect the intricate non-linear relationships that may exist among specific disease markers, chromatin accessibility sites, and transcription factors. Building on this, scAB [10] extended the method by applying matrix factorization to the same Pearson correlation matrix and assigning greater weights to bulk samples displaying the phenotype of interest. This was achieved through the imposition of a penalizing diagonal matrix that represented the likelihood of a sample manifesting a disease phenotype. Later on, the PACSI model [8] calculated the correlation scores between cells and phenotypes by summarizing the distances between SC and patient sample gene signature defined modules within the protein-protein interaction network. Despite achieving significant performance across a variety of SC and spatial transcriptomic (ST) datasets in identifying disease-associated subgroups of cells and tissues, these methods have one drawback: they assessed the similarity between SC and bulk/patient samples on an individual basis and failed to capture the collective statistical characteristics of SC groups and patient samples.

The DEGAS framework [3] enhanced prior methodologies by aligning the statistical distributions of SCs and patients within a unified latent feature space. Utilizing a multi-task transfer learning (TL) neural network (NN), it facilitated the alignment of SC and patient distributions by minimizing the maximum mean discrepancy (MMD) between their features. This process enabled the direct transference of patient-level attributes onto SC samples. The model has been demonstrated to be a powerful tool for identifying disease-related subgroups of SCs in conditions such as Alzheimer’s disease (AD) and multiple myeloma (MM), and has been successfully applied to prostate cancer ST datasets [12]. Building on the success of the DEGAS framework, we enhanced it by incorporating the Wasserstein loss function to quantify the distribution differences between SC/ST and patient samples. This function is a well-established probability metric derived from optimal transport (OT) theory and exhibits superior geometric and statistical properties compared with the MMD loss [13]. Additionally, in this updated version of the DEGAS model (DEGAS v2), we have integrated additional loss function options tailored for time-to-event prediction tasks. Similar to the original DEGAS framework, our method includes three steps: 1) preprocessing SC/ST and bulk/patient samples to identify genes reflecting cellular heterogeneity and disease trait variations, ensuring both sample types share the same feature/gene set for input into the NN; 2) training a multi-task TL deep-learning (DL) framework, where SC/ST and bulk/patient samples are embedded in the same feature space by minimizing their Wasserstein distance; and 3) evaluating SC/ST samples by applying the patient attributes prediction NN to SC/ST features and extracting their disease-associated scores.

To demonstrate our framework’s superiority, we first conducted a comprehensive benchmark study on three prostate cancer ST datasets (10X Genomics; [14]), comparing our extended DEGAS framework (DEGAS v2) with other SOTA methods including Scissor [9], scAB [10], and PACSI [8]. Then, we replicated the AD study in [3], under-scoring our model’s robustness regarding the selection of disease indices, which reflect the same disease progression from different perspectives. Lastly, we carried out an exploratory study on the multiple myeloma dataset, enabling the discovery of SC subgroups exhibiting a high hazard for disease progression and substantial differences in drug response during different time frames. This paper is structured as follows: Section 2 delineates the explicit form of the TL problem, the Wasserstein-based model for TL, and our specialized loss functions. We then expound on the experimental settings for various ST and SC datasets and define the precise aims of each experiment. Section 3 will illustrate and discuss the findings from three ST prostate cancer datasets, two SC AD datasets, and the MM survival and drug response discovery dataset. Finally, Section 4 will engage in a discourse on the merits and limitations of our proposed framework.

## METHODS

### DEGAS Framework

#### Overview

To investigate the disease association of SC/ST datasets, the DEGAS v2 model employed a TL framework to associate SC/ST samples with bulk/patient instances and assign patient-level disease attributes to SC/ST samples regarding their feature-level similarity. For model inputs, we denoted the preprocessed SC/ST and bulk/patient GEM by *X*_*SC*_ ∈ ℝ^*NSC*C/*ST*,*Ngenes*^ and *X*_*pat*_ ∈ ℝ^*Npat*,*Ngenes*^, respectively. Each column of these two gene matrices corresponded to the same gene. For SC-level model outputs at the training phase, we incorporated cell types or tissue histology annotations, if there were available, as SC/ST prediction targets and denoted them by *y*_*SC*_ ∈ ℝ^*SC*C/*ST*^. For the bulk/patient-level outputs, we assigned them binary labels: *y*_*pat*_ ∈ ℝ^*Npat*^ to indicate high risk (i.e., low drug response) and low risk (i.e., high drug sensitivity), respectively. We also allowed the analysis of the time-to-event data, where *t*_*pat*_, *c*_*pat*_ ∈ ℝ^*Npat*^ signified event time and status (“1” for event of interest and “0” for censoring). During the evaluation phase, our goal was to associate each SC with patient-level binary classifi- cation softmax score or survival hazard score. The model framework was summarized in Fig. 1.

**Fig. 1.**
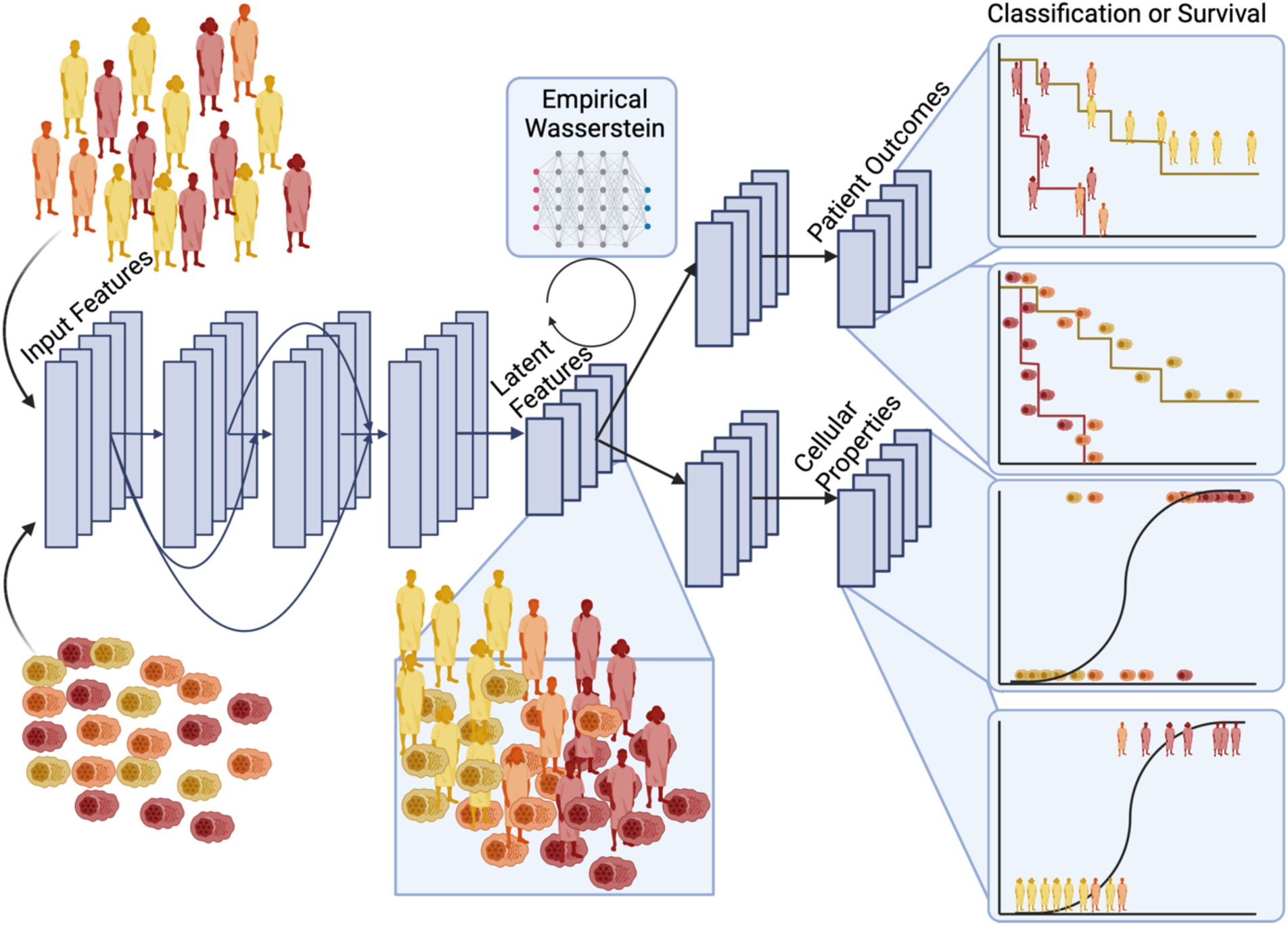
Overview of DEGASv2 Framework

#### DEGAS network architecture

The DEGAS v2 model comprised of four NN components: a feature extractor shared by SC/ST and bulk/patient samples, a discriminator designed for applying the OT based-algorithm to distinguish SC/ST- and bulk/patient-level features, and two classifiers built for domain specific tasks, i.e. stratifying high and low risk patients, and classifying SC/ST samples as different cell types/tissue classes (**Fig. 1**). DEGAS v2 aligns bulk/patient-level disease attributes to SC/ST samples through a two-step process: training and evaluation.

During the training stage, SC/ST inputs *X*_*SC*/*ST*_ and bulk/patient input *X*_*pat*_ were embedded into a common latent feature space using a shared feature extractor function, *f*_*extractor*_. Inspired by the concept of ResNet [15], *f*_*extractor*_ was implemented as a three-layer dense NN, which combined features extracted from shallower layers (low-level features) with those from the current layer as the input for the next layer. Our approach employed skip connections to enhance the complexity and the diversity of the feature space. The *f*_*extractor*_ is defined as,

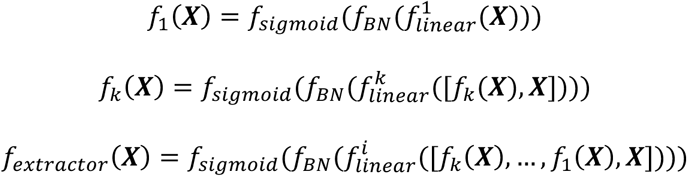

In the above function, *f*^*i*^_*linear*_ is a linear function defined by *f*^*i*^_*linear*_ = 𝑾^𝑖^𝒙 + 𝒃^𝑖^, where 𝒙 ∈ ℝ^*N*𝑖^, 𝑾^𝑖^ ∈ ℝ^𝑀𝑖,*N*𝑖^, 𝒃 ∈ ℝ^𝑀𝑖^, with *N*_𝑖_, and 𝑀_𝑖_ input and output features respectively. Each linear layer is incorporated with a dropout regularization function (p = 0.5), a 1d batch normalization function (*f*_𝐵*N*_), as well as a Sigmoid activation function (*f*_𝑠𝑖2𝑚𝑜𝑖𝑑_) to scale non-linear features. For model reproducibility, we initialized the model weights 𝑾^𝑖^ using the uniform distribution within range [0, 1] with fixed random seeds. Biases were initialized as 0 for all layers.

#### Transfer learning loss function

After embedding bulk/patient-level and SC/ST into the same feature space, we required a metric to quantify the discrepancy between *f*_*extractor*_(*X*_*SC*_) and *f*_*extractor*_7*X*_*pat*_8. Traditionally, the kernelized MMD loss function is introduced for its sensitivity in detecting distributional discrepancies [16–18]. However, as previously shown [13] highlighted the computational intensity of the MMD, which limited its use in large-scale SC datasets. Furthermore, they observed that specific kernels can cause the MMD distance to lack regularity, including continuity and differentiability. These limitations hinder our ability to use the kernelized MMD as an ideal measurement of distributional differences between SC/ST and bulk/patient features.

Recognizing this limitation, we introduced the Wasserstein loss function, which was inspired by the Kantorovich-Rubinstein duality of the OT problem [19] and widely adopted in numerous generative models and TL tasks. The Wasserstein loss was considered as a well-defined metric between distributes and demonstrated its superiority over other metrics in terms of its stability and scalability [13, 20, 21]. Specifically, the Wasserstein-1 distance between distributions on two feature domains is defined as follows.

**Definition 1** (Wasserstein-1 Distance). *Let* ℙ^𝑠^ *and* ℙ^𝑡^ *be the probability distribution of features from the source (bulk/patient) and the target (SC/ST) domains, respectively, their Wasserstein-1 distance reads,*

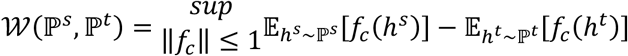

*where* ℎ^𝑠^ *and* ℎ^𝑡^ *correspond to the source (bulk/patient) and target domain (SC/ST) samples/features, respectively. The supremum is taken over all 1-Lipschitz functions f*_𝑐_*, which could be realized using a multi-layer NN. In our framework, it was designed as a three-layer fully connected NN with output dimensions of 12, 12, and 1, respectively. Each linear layer was incorporated with a dropout function (p = 0.5) a batch normalization (BatchNorm1d) function, and LeakyReLU (α = 0.2) activation function*.

In each training iteration, we randomly balance sampled SC/ST and bulk/patient instances with batch size 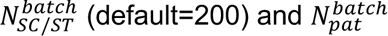 (default=200) (default=200), respectively. Then the empirical version of Wasserstein distance was reformulated as the following where 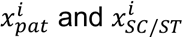 were a minibatch of bulk/patient and SC/ST samples respectively.

**Definition 2** (Empirical Wasserstein-1 Distance)

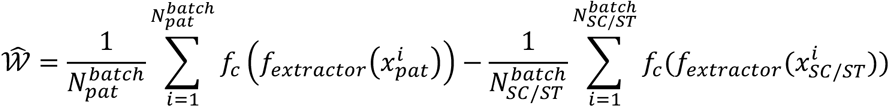

The interpretation of this target function is similar to that of traditional generative adversarial models. Our goal is to create domain-invariant features so that the discriminator function *f*_𝑐_ couldn’t differentiate between them. Therefore, we updated *f*_*extractor*_ and *f*_𝑐_ alternatively with opposite purposes: *f*_*extractor*_ was optimized to minimize the target function while *f*_𝑐_ was fine- tuned to maximize it. Since *f*_*extractor*_ is involved in other loss functions, we will give its optimization target function later.

The target function for optimizing *f*_𝑐_ is defined as:

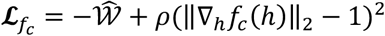

where the second term (‖∇_ℎ_*f*_𝑐_(ℎ)‖_2_ − 1)^2^ is a gradient penalty function, designing to ensure that *f*_𝑐_ satisfies the 1-Lipschitz constraint [22]; its input vector ℎ^𝑖^ was synthesized by interpolation between 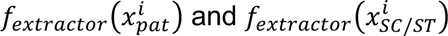 Specifically, we generated a number τ uniformly within [0,1] and 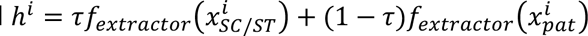. The weight 𝜌 was default set to be 10.

#### Task specific loss functions

As mentioned earlier, *f*_*extractor*_ was designed for providing informative features for our target task, i.e. distinguishing high and low hazard SC/ST samples. To achieve this goal, *f*_*extractor*_should have two properties: the ability to represent the main characteristics of each cluster of cell types, while simultaneously being able to distinguish high/low hazard subgroups of cells within each cell cluster. Therefore, our task-specific target functions consist of the classification loss function for SC samples and the hazard classification/prediction loss function for bulk/patient samples. The SC prediction loss is defined below.

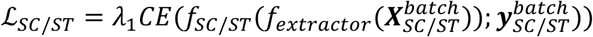

The variable *f*_*SC*/*ST*_ is a one-layer dense NN with an input dimension N^3^ and an output dimension 1. CE represents the cross-entropy loss function for multi-class classification. In the default setting, we set λ1 = 1.0, but when SC/ST labels are infeasible, it was set to be 0.

Similarly, two other options of loss functions for patient-level prediction are included in our framework.

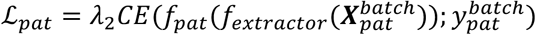

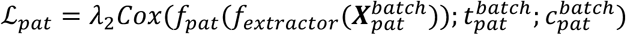

The first loss function was formulated for binary classification tasks, such as identifying the presence or absence of a disease or drug response. The second loss function was designed for time-event outcomes, with 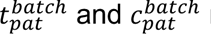 representing the time and event labels (“1” event, “0” censor), respectively. For binary classification, *f*_*pat*_ is defined as a simple linear layer with an input dimension N^3^ and an output dimension 1. For survival prediction tasks, *f*_*pat*_ is defined as a linear layer with the same input and output dimensions, combined with a dropout function (dropout rate = 0.5), a batch normalization (BatchNorm1d) function, and a Sigmoid activation function. The notations CE and Cox correspond to the cross-entropy and the Cox proportional hazard loss [23], respectively. Combining the Wasserstein loss function with the task-specific loss function yields the following target function for updating *f*_*extractor*_, *f*_*SC*/*ST*_, and *f*_*pat*_ simultaneously.

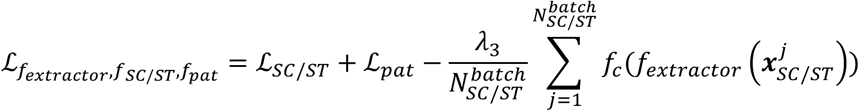

In the above equation, we include the second term from **Eq. 3** by referring [13].

#### Model optimization

We grouped *f*_*extractor*_, *f*_*SC*/*ST*_, and *f*_*pat*_ together into the same loss function separate from the *f*_𝑐_ loss function. The model was optimized by iteratively minimizing these equations in an alternating pattern. During this process, we backpropagated weight gradients using the Adam optimizer [24], with the learning rate, beta parameters, and training iterations set to be 0.01, (0.9, 0.999), and 300, respectively. Unlike conventional Wasserstein loss-based TL frameworks [20, 21], we updated the discriminator *f*_𝑐_ less frequently than the *f*_*extractor*_ feature extractor to prevent SC features from converging too closely to patient features, thereby preserving cell-type specific characteristics. This approach also offered greater flexibility compared to the MMD loss, as it allowed us to regulate the proximity between SC and patient features through the update frequencies of the feature extractor and discriminator. Under the default setting, we updated *f*_𝑐_ every other iteration. When updating *f*_𝑐_, we detached all features *f*_*extractor*_(*X*^𝑏𝑎𝑡𝑐ℎ^) and *f*_*extractor*_(*X*^𝑏𝑎𝑡𝑐ℎ^) from the computational graph so that *f*_*extractor*_ wouldn’t be affected by this step. Similarly, parameters of *f*_𝑐_ were fixed when optimizing *f*_*extractor*_, *f*_*SC*/*ST*_, and *f*_*pat*_.

We applied Bootstrapping Aggregation (Bagging) [25] to improve the model stability. For each sub-model and each training iteration, we randomly sampled 200 bulk/patients and 200 SC/ST instances using distinct and pre-defined random seed. Additionally, we improved the model diversity by initializing NN weights using different pre-defined random seeds for different sub-models, leading to distinct initial points for optimization.

#### Model evaluation

During the evaluation, the patient-level disease attribute predictor *f*_*pat*_ was utilized on SC/ST features *f*_*extractor*_(*X*_*SC*/*ST*_) to assign hazard scores to SC/ST samples. To derive normalized hazard scores ranging between [0, 1], we implemented distinct methods for categorical and time-to-event prediction outcomes. Specifically, for binary classification tasks, we employed the Softmax function to convert the predicted scores to values within the [0, 1] range and extracted probability scores for the high hazard (labeled as “1”) class as hazard scores. In the case of time-to-event predictions, since the outputs was processed through the Sigmoid function, they naturally yielded normalized scores within this range. Ultimately, the averaged normalized scores from various sub-models served as the final prediction score for each SC/ST sample.

#### Datasets

We conducted comprehensive experiments and benchmark studies on a wide range of ST and SC datasets and used ground truth (GT) labels for evaluation and comparison. The ST experiments consisted three datasets: Erickson patient 1 [14], Erickson patient 2 [14], and the10x Genomics. Patient-level samples were collected from the TCGA PRAD cohort [11], and patients were associated with time-to-event labels, based on their progression-free survival (PFS). Further details will be provided in the Spatial Transcriptomics Experiments section.

For the SC experiments, we included two AD SC datasets: the Grubman [26] and the Mathys [7], with each SC labeled by its cell type. The Grubman dataset included nuclei from the entorhinal cortex tissue of 12 individuals with AD and normal controls (NC) from the Victorian Brain Bank cohort [26]. Researchers then refined this dataset to 13,214 cells, each characterized by 10,850 genes by applying quality control (QC) procedures. They finally identified 8 cell types including microglia (mg), astrocytes (astro), neurons, oligodendrocytes (oligo), oligodendrocyte progenitor cells (OPCs), endothelial cells (endo), unidentified cells (unID), and dou- blet cells (doublet). The Mathys SC dataset was collected from the prefrontal cortex of 48 individuals participating in the Religious Orders Study (ROSMAP) cohort [27].

Among these individuals, half exhibited AD pathology, while the remainder showed no pathology. Researchers then refined it to 70,634 cell nuclei with 17,926 genes after QC. Through clustering and investigating statistical enrichment of marker gene sets, they identified 8 cell types including excitatory neurons (Ex), inhibitory neurons (In), astrocytes (Ast), oligodendrocytes (Oli), microglia (Mic), oligodendrocyte progenitor cells (OPC), endothelial cells (End), and pericytes (Per). For patient-level inputs, we utilized the MSBB [28] RNA-seq data, which comprised samples from 135 AD patients and 86 normal control individuals. The ages of participants ranged from 61 to over 90 years.

#### Data Cleaning and Preprocessing

We designed convenient built-in functions to standardize the data preprocessing pipeline based on the Seurat package [29]. For SC/ST data, we utilized the “createSeuratObject()” function to transform SC and ST gene count matrices into Seurat objects. This step involved the exclusion of genes expressed in an insufficient number of cells/ST samples (min.cell = 400) and the removal of cells/ST spots with insufficient gene counts (min.features = 200). To further ensure data quality, we employed the ”subset()” function to filter out SC/ST samples containing an excess of mitochondrial genes or a deficit in gene counts, using the criteria percent.mt ¿ 0.95 and nFeature RNA ¡ 0.05, respectively.

To develop feature sets from DEGs in the ST and SC datasets, we followed standard Seurat clustering processes: Log transformation using ‘NormalizeData(scale.factor = 10000)‘, data scaling through ‘ScaleData()‘, PCA analysis via ‘RunPCA()‘, construc- tion of a KNN graph using ‘FindNeighbors()‘, and identification of sample clusters with ‘FindClusters()‘. Subsequently, we used the ‘FindAllMarkers‘ function (parameters: only.pos = FALSE, min.pct = 0.25, logfc.threshold = 0.25) to extract DEGs for each cluster. Within each cluster, genes were then selected based on the average log- fold change, retaining only the top 10% of genes.

Furthermore, a filtering step based on p-values was applied, again preserving the top 10% of genes.

For bulk/patient-level data, individuals with absent clinical follow-up information were removed. Then, we removed duplicated gene names by adding their counts together. Only common genes to both the bulk/patient and SC/ST datasets were retained. To understand how feature selection influenced model performance, we developed three feature sets using different approaches: (1) The top 250 genes exhibiting the highest variance among patients. (2) DEGs identified in the SC/ST preprocessing phase. If the number of candidate genes exceeded 250, we intersected this candidate gene set with the top 500 highest variance genes among patients to reduce the feature set size. To get a smaller candidate gene set, we could shrink the highest variance gene set of patients from top 500 to top 250 genes with the step size 50. (3) Most significantly DEGs between patient groups stratified by median PFS time.

After the feature selection process, we rescaled both the SC/ST and bulk/patient gene count matrices by applying a transformation of 1.5^log2(counts+1)^ to all raw counts. Then, within each sample, we calculated the z-score for each gene by subtracting the mean of the transformed gene counts and then dividing the result by their standard deviation. Finally, for each gene over all samples, we scaled the z-score into [0, 1] by min-max scaling. For ST samples, we further concatenated all processed ST slides into a large matrix by stacking their samples. After all of these preprocessing steps, we had a SC /ST data matrices: *X*_*SC*/*ST*_, the associated SC/ST labels (if available) *y*_*SC*/*ST*_, the bulk/patient data matrix *X*_*pat*_, and its associated labels *y*_*pat*_ or *t*_*pat*_ (time) and *c*_*pat*_ (censor).

#### Spatial Transcriptomics Experiments

The ST experiments encompassed three distinct prostate ST cohorts, namely, the 10x Genomics, the Erickson patient 1 [14], and the Erickson patient 2 [14] datasets. The 10x Genomics dataset comprised four distinct slides, representing samples from normal, stage II, stage III, and stage IV patients. The staging information for the 10x Genomics data facilitated an evaluation of the DEGAS model’s ability to rank overall hazard scores for ST slides from the different patients: the intuition being that increasing stage patients should have biopsies with increasing overall DEGAS risk scores. On the Erickson patient 1 dataset, each ST spot was assigned a histology annotation by the original authors, serving as GT labels for model evaluation and benchmark studies. The Erickson patient 1 dataset was used to understand how DEGAS predictions aligned with histopathological tissue labels. The Erickson patient 2 dataset lacked GT histopathological labels and served as an exploratory dataset. We employed a specific parameter setting for training on all ST datasets: λ1 = 0.0, λ2 = λ3 = 3.0, with other parameters set to default values. For the 10x Genomics dataset, we vertically concatenated samples from all four stages into a large data matrix, facilitating the evaluation of the model’s ability to discern differences in hazard scores across various stages. The model was run separately for the Erickson Patient 1 and Patient 2 dataset, as the task was not to compare risks scores between samples. For each cohort, DEGs of ST samples (feature set) were utilized as the input feature set.

On the 10x Genomics dataset, our approach incorporated the Wilcox nonparametric test between DEGAS hazard scores across varying cancer stages. It illustrated the DEGAS model’s proficiency in differentiating the severity of cancer across stages. For model evaluation on the Erickson Patient 1 dataset, we employed the tumor versus non-tumor binary classification task to assess the model performance. Specifically, we labeled the GG1, GG2, GG4, and GG4 Cribriform annotated regions as tumor areas, with all other regions classified as non-tumor. We then calculated the ROCAUC score of DEGAS hazard scores for this binary tumor classification task. It’s worth noting that we didn’t include any human annotations in training ST samples.

Thus, the results were purely inferred from the correlation between ST and patient samples. Besides, boxplots of hazard scores in each annotated region were depicted to indicate the association between hazard scores and tissue types. Regarding the Erickson Patient 2 dataset, our current methodology focused on the spatial visualization of hazard scores and left it as our further work to investigate biological processes potentially revealed by DEGAS scores.

We compared the DEGAS model with three state-of-the-art (SOTA) models: Scissor [9], scAB [10], and PACSI [8]. Originally designed for identifying high-risk SC samples, these models could also be applied to ST datasets without significant challenges. We first assessed the efficacy of these models in identifying tumor-related high-risk regions using the annotated Erickson Patient 1 dataset. Given that the bench- mark methods provided categorical predictions, the computation of ROCAUC scores was not feasible for performance measurement. Consequently, we employed bar plots to visually represent the proportion of prediction categories within both tumor and non- tumor regions. The prevalence of high hazard regions within tumor areas acted as a surrogate for the recall score in model comparison. To identify consistency between the DEGAS model predictions and other methods, DEGAS hazard scores were visualized as boxplots and grouped based on their Scissor, scAB, and PACSI labels. Non-parametric Wilcox tests were then performed to test whether Scissor Positive regions exhibited higher DEGAS hazard scores than other regions. Similar procedures were performed for the scAB and the PACSI models. Finally, for model comparison on the 10x Genomics dataset, we demonstrated the proportion of high-hazard region selected by benchmark methods for slides from different cancer stages.

#### Alzheimer’s Disease Experiments

We assessed our model’s performance on two AD SC datasets: the Grubman (SC) [26], the Mathys (SC) [7]. For patient-level inputs, we employed the MSBB [28] datasets. Notably, the MSBB dataset comprises comprehensive clinical measurements of patients such as Braak stages, CERAD scores, Clinical Dementia Ratings (CDR), and plaque mean scores. To accommodate these scoring systems with our binary classification setting, we stratified patients into high and low-risk groups with following criterions: CERAD score of 2 or less were classified as high risk. A Braak stage of 5 and above, CDR score of 3 and above, was also indicated high risk. For the plaque mean score, we excluded patients with missing values and used the median score as a threshold for risk categorization. After applying the QC, feature selection, and preprocessing steps, we employed the DEGAS model to infer the above four different binarized measurements one-by-one to the Grubman and the Mathys SC samples, respectively (parameter settings: λ1 = 1.0, λ2 = λ3 = 3.0.

To validate our hypothesis that the DEGAS model was able to differentiate cells from AD and NC patients, we stratified SCs into distinct groups based on their cell and the disease label of the patients. Intuitively, we expected that cells sampled from AD patients should exhibit in general higher disease associated scores than cells from NC individuals. To verify our assumption, we employed a non-parametric Wilcoxon test to compare DEGAS hazard scores of cells originating from AD and NC patients across all cell types and all disease attributes.

Additionally, we examined the congruence between various scoring systems by assessing the correlation of DEGAS hazard scores, derived from different disease attributes such as CDR and plaque mean scores. Furthermore, we studied cells with inconsistent CDR and Braak scores such that the cell has an atypical, inferred phenotype. We implemented a linear regression analysis for each cell type across all scoring pairs, e.g. CDR versus plaque mean scores.

Outliers were identified by selecting cells corresponding to residuals with absolute values exceeding 2σ, the standard deviation of all residuals from the linear regression. For instance, oligodendrocytes cells with residuals (CDR as response variable and plaque mean score as predictor variable) larger than 2σ were classified as having high CDR and low plaque mean scores. By employing the “FindMarkers()” function of the Seurat package on these outlier cells, alongside cells with residual absolute values smaller than 2σ, we successfully identified a list of genes that are potential markers for these distinct outlier cell subpopulations.

#### Multiple Myeloma Experiments

The primary objective of the multiple myeloma (MM) experiment was to identify cells with a strong correlation to malignant clinical outcomes. This experiment was conducted using SCs from 49 myeloma patients, as documented in the Indiana University School of Medicine (IUSM) MM dataset [30]. Specifically, 325,025 high-quality CD138+ myeloma cells were utilized, having been meticulously selected to exclude low-quality and non-plasma cells. These cells were then stratified into 25 clusters using Seurat, each cluster representing a unique transcriptomic profile. At the patient level, transcripts per million (TPM) values of bulk tissue RNA profiles, along with their corresponding PFS times and event/censor indicators, were collected from the MMRF. The MMRF also provided longitudinal data from which the anytime response and extended responses to proteasome inhibitor and immunomodulatory drug therapy could be evaluated. This expansive patient cohort broadened our goal to ascertain whether, at the cellular level, survival-related hazard scores are consistent with drug responses. Another vital goal is to determine whether immediate or extended drug responses are more closely associated with disease progression post-treatment.

For predicting survival hazard scores, we implemented the standard feature selection and preprocessing pipeline. To obtain a more concise set of gene candidates, we retained DEGs between high-risk individuals (patients with a PFS event within 3 years) and low-risk individuals (other patients) by setting the p-value threshold to be 0.001 (default = 0.05). For predictive outcomes, instead of using the time-to-event labels, we chose the binarized survival label from the previous step and yielded more stable results during training. Subsequently, we used the “ClassClass” model along with the Wasserstein loss function to train 10 submodels, allocating 300 training iterations for each. The NN’s output was normalized by a softmax function to yield a scaled PFS hazard score. To highlight the differences among various SC groups during model training, we set lambda1 at 3.0 and conducted 500 training iterations. For the extended response prediction, we utilized identical input gene matrices but limited the training to 100 iterations. These hyperparameters were selected empirically to underscore the distinctions among the different cell groups.

## RESULTS

### DEGAS identifies high risk regions in Prostate Cancer ST

We presented an in-depth investigation of the DEGAS model’s performance on ST datasets, where the availability of GT histological labels facilitated rigorous model evaluation. We first confirmed the DEGAS model’s ability to rank disease severity and identify disease associated tissues using two prostate ST cancer datasets: Erickson patient 1 [14], and the public 10x Genomics datasets. The Erickson patient 1 and 10x Genomics datasets were associated with pathologists’ annotations and clinical information, respectively. The Erickson patient 2 dataset was used as the exploration dataset. Additionally, we conducted a comprehensive benchmark study by comparing DEGAS against three SOTA models: Scissor [9], scAB [10], and PACSI [8]. Our research findings underscored the DEGAS model’s superior ability in detecting tumor and disease-associated tumor microenvironment (TME) regions within ST slides and ranking hazard scores for tissues from different risk stages.

Using PFS as the prediction target, the DEGAS model exhibited a remarkable ability to accurately assign disease-related severity scores to different slides and regions. On the 10x Genomics dataset, the average DEGAS score increased steadily as the cancer staging ranged from normal to stage IV (**Fig. 2A**) and the proportion of Scissor positive cells also increased with stage (**Fig. 2B**). The DEGAS association also tended to be greater in Scissor positive cells in comparison to Scissor negative cells representing high concordance between DEGAS and Scissor in the 10X Genomics dataset (**Fig. 2C**). P-values of the Wilcox non-parametric test of DEGAS scores between difference stages were significant, underscoring the potential utility of DEGAS as a diagnostic tool for identifying the tumor stage of patients (**Fig. 2C**)

**Fig. 2.**
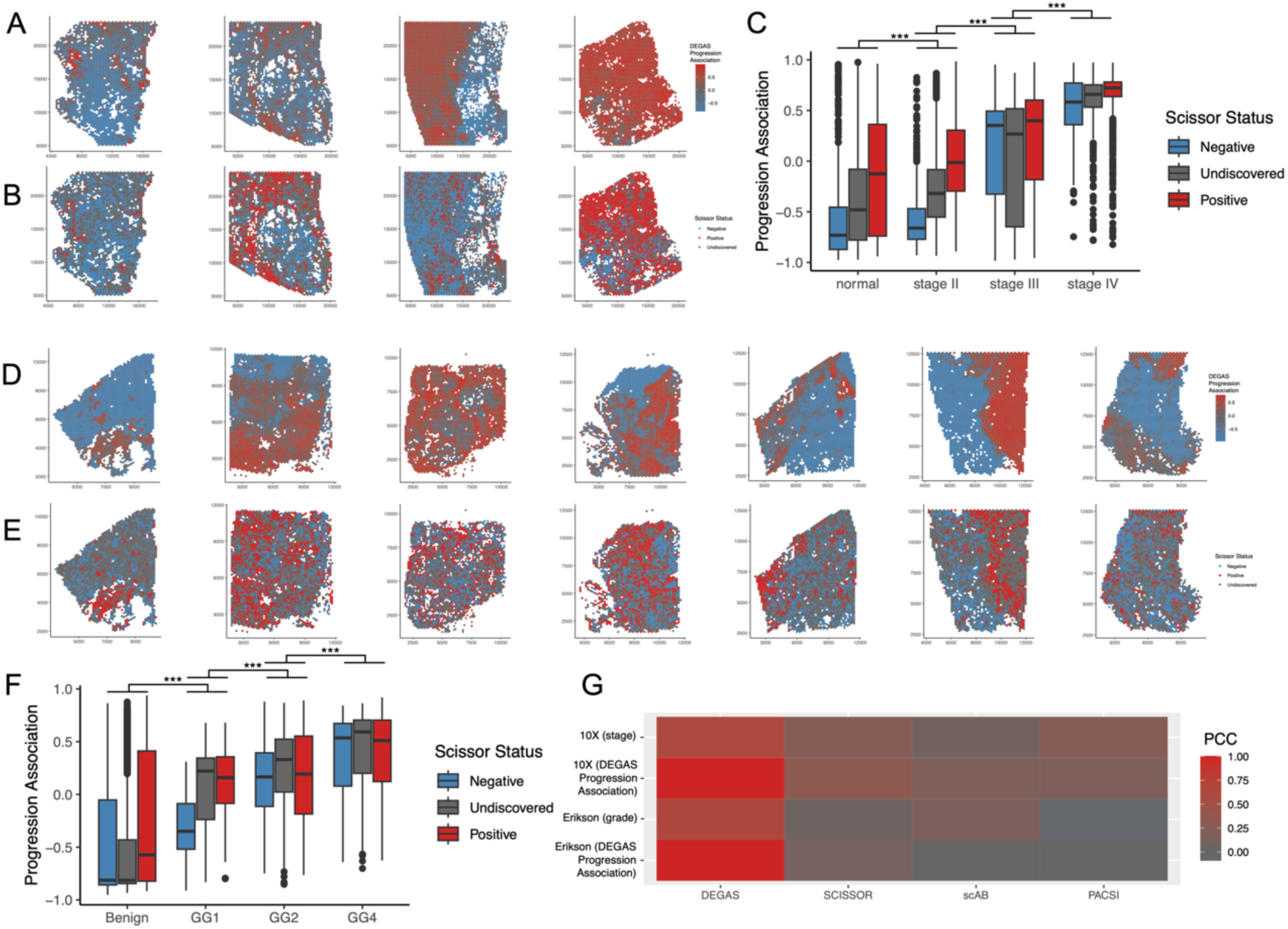
DEGAS Version 2 performance on prostate cancer spatial transcriptomics data. **A)** DEGAS Progression association and **B)** Scissor predictions on 10X Genomics samples. **C)** DEGAS Progression association and Scissor predictions across prostate cancer stage in 10X Genomics samples. **D)** DEGAS Progression association and **E)** Scissor predictions on Erikson et al. samples. **F)** DEGAS Progression association and Scissor predictions across prostate cancer stage in Erikson et al. samples. **G)** Comparison of DEGAS, Scissor, scAB, and PACSI to ground truth grade/stage and to DEGAS progression associations.

On the Erickson patient 1 dataset, DEGAS scores again aligned tumor tissue specificity (**Fig. 2D**) and was concordant with Scissor positive regions (**Fig. 2E**). DEGAS scores also increased significantly with the tumor grade (**Fig. 2F**). DEGAS progression associations were more correlated to stage (10X Genomics) and grade (Erickson) than Scissor, scAB, or PACSI outputs (**Fig. 2G**). It is worth noting that all of the methods had positive correlations with stage and grade except for a slightly negative PACSI correlation with grade (**Fig. 2G**).

### DEGAS Identifies cells with AD-associated and Asymptomatic AD-associated phenotypes

To evaluate the performance of DEGAS and identify AD-associated subtypes of cells, Braak score, CDR, CERAD, and Aβ plaque associations were overlaid onto the cells from the Grubman et al. scRNA-seq dataset. We found that 7/8 of the cell types exhibited greater Braak- associations (**Fig. 3A**), 6/8 of the cell types exhibited greater CDR-associations (**Fig. 3B**), 5/8 of the cell types exhibited greater CERAD-associations (**Fig. 3C**), and 7/8 of the cell types exhibited greater Aβ plaque-associations (**Fig. 3D**) in cells originating from AD patients in comparison to CN patients. These cell types formed distinct clusters with heterogeneity between cells derived from AD and CN patients (**Fig. 3E,F**). The overall AD-associations (i.e., Braak, CDR, CERAD, and Aβ plaque combined), also tended to correlate with the AD status of the patients (**Fig. 3G,H**). Notably there were cells from AD patients that were not AD-associated and cells from CN patients that were AD-associated (**Fig. 3G,H**). All of the AD association scores were correlated with cell type and AD metric dictating the strength of the correlation (**Fig. 3I**). In the baseline comparisons between DEGAS, Scissor, scAB, and PACSI, we found that DEGAS had much more significant differences in score between AD and CN samples regardless of cell type (**Fig. 3J**).

**Fig. 3.**
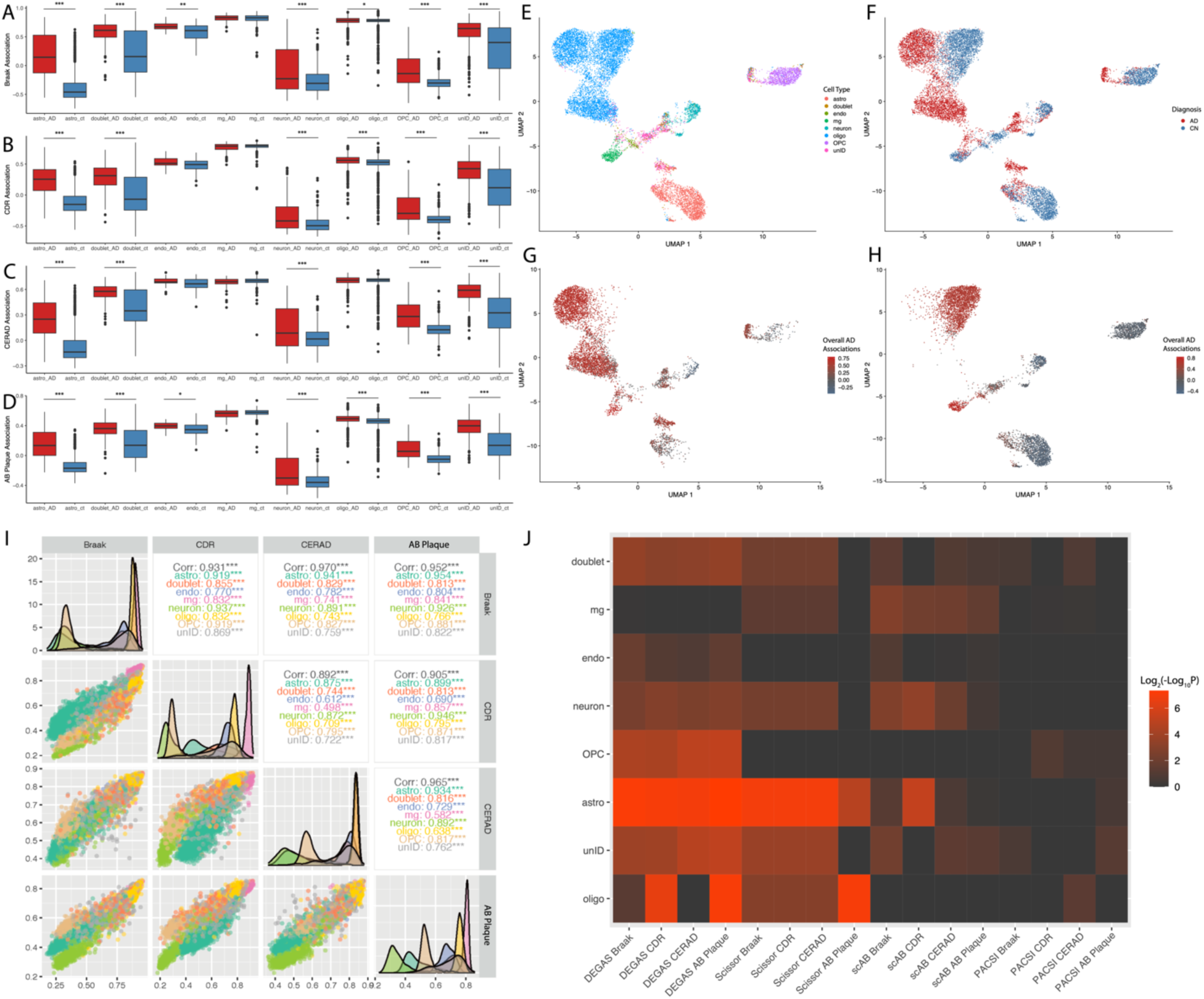
DEGAS Version 2 performance on AD single cell transcriptomics. Comparison of Braak **(A)**, CDR **(B)**, CERAD **(C)**, and AB plaque **(D)** DEGAS associations between cells from AD and CN samples. **E)** Cell types in Grubman et al. dataset. **F)** AD status of the samples for each cell in the Grubman dataset. **G)** Overall AD associations (Braak + CDR + CERAD + AB plaque) in cells from AD patients. **H)** Overall AD associations from in cells from CN patients. **I)** Correlation of different AD association scores across different cell types. **J)** Performance of different cell prioritization tools to correctly predict cells from AD patients as having higher AD association based on -Log10P.

### DEGAS Identifies Sham Response Myeloma Cells at High Risk of Progression

Finally, we explored the effectiveness of DEGASv2 in predicting survival-associated hazard and drug response metrics in myeloma scRNA-seq. Based on our previous work, the 325,025 cells clustered into 25 clusters (**Fig. 4A**). Some of these clusters had associations with disease stage (**Fig. 4B**) and proliferation (**Fig. 4C**). Clusters 11, 20, and 22 had significantly greater DEGAS progression associations than the other clusters (**Fig. 4D,E**) and expressed increased levels of *PHF19* (**Fig. 4F**)[30]. Interestingly, clusters 11 and 20 also had significantly greater associations with anytime response to PI + IMiD therapy (**Fig. 4G,H**). However, these same clusters, 11 and 20, had significantly lower associations with extended response to PI + IMiD therapy, i.e., greater than 60 days (**Fig. 4I,J**). Though paradoxical, clusters 11 and 20 may represent cells that have an initial response but eventually evade therapy. We therefore evaluated the differences between the progression associated clusters 11, 20, and 22. Based on this analysis we identified multiple DEGs specific to cluster 11 (**Fig. 4K**), cluster 20 (**Fig. 4L**), and cluster 22 (**Fig. M**). The top markers for cluster 11 were FAM111B (**Fig. 4N**) and WDR76 (**Fig. 4O**).

**Fig. 4.**
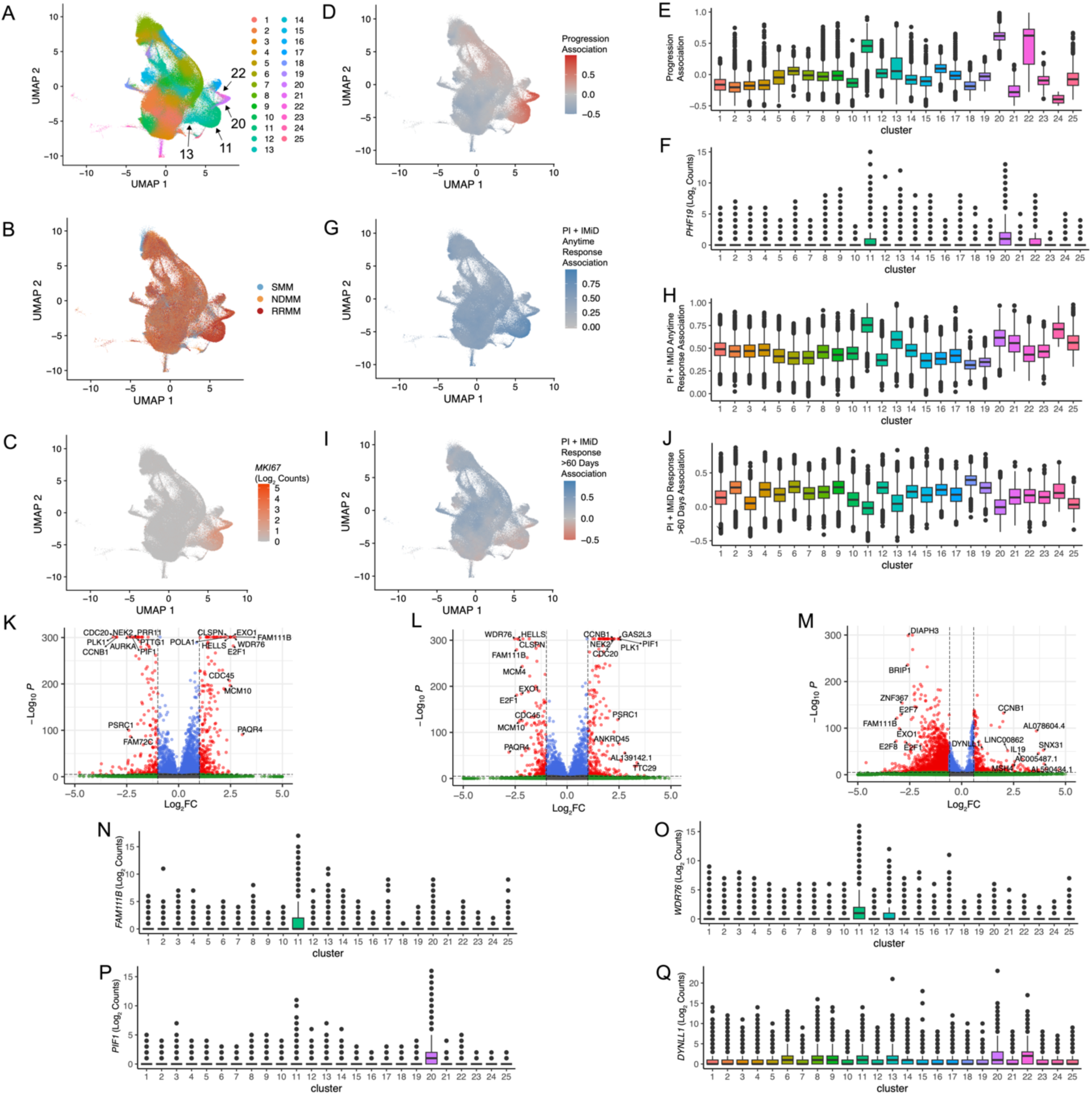
Exploratory analysis of MM data with DEGASv2. **A)** Clusters analysis of myeloma cells. **B)** Diagnosis of patient samples from which cells were derived. **C)** *MKI67* expression in cells. **D)** DEGAS Progression Association overlaid onto cells with boxplots to compare relative values between clusters **(E)**. **F)** *PHF19* expression of cells displayed across different clusters. **G)** DEGAS Anytime PI + IMiD response association overlaid onto cells with boxplots to compare relative values between clusters **(H)**. **I)** DEGAS Extended PI + IMiD response (>60 days) association overlaid onto cells with boxplots to compare relative values between clusters **(J)**. **K)** Differential expression of cluster 11 compared to cluster 20 and 22. **L)** Differential expression of cluster 20 compared to cluster 11 and 22. **M)** Differential expression of cluster 22 compared to cluster 11 and 20. Expression of novel marker genes for clusters 11, 20, and 22 namely: *FAM111B* **(N)**, *WDR76* **(O)**, *PIF1* **(P)**, and *DYNLL1* **(Q)**.

Interestingly WDR76 was also upregulated in cluster 13 (**Fig. 4O**). The top marker for cluster 20 was PIF1 (**Fig. 4P**) and for cluster 22 was DYNLL1 (**Fig. 4Q**). Cluster 22 the progression associated cluster without the short response transcriptional phenotype had DIAPH3, BRIP3, ZNF367, E2F7, FAM111B, EXO1, E2F8, and E2F1 as the top 10 downregulated genes in comparison to the clusters (11 and 20) with the short response transcriptional phenotype (**Fig. 4M**). This gene set was significantly enriched for REACTOME TP53 REGULATES TRANSCRIPTION OF GENES INVOLVED IN G1 CELL CYCLE ARREST (Reactome: M27635, FDR = 1.78•10^-6^) possibly indicating a relationship between these proliferative cells and *TP53* alterations in these resistant cells. Another notable cluster 11 marker related to chemotherapy resistance was PAQR4 (**Fig. 4K**). There were also multiple proliferation related markers that differed between these clusters including NEK2, CDC20, CCNB1, and MCM10 (**Fig. 4 K-M**).

## DISCUSSION

The inference of disease attributes at the SC level presents a promising avenue for elucidating underlying disease biological processes, monitoring drug responses, and prioritizing drug targets. Given the lack of SC-level disease-related annotations, the strategy of transferring “impressions” of patient-level disease attributes to each SC offers an attractive approach to discovering SC sub-populations of interest. Building upon the existing DEGAS framework, which aims tackle this challenge, we extended it by replace the original kernelized MMD loss function by a computationally and theoretically more efficient Wasserstein loss. This enhancement led to superior performances in aligning patients’ and SCs” gene expression values. Furthermore, we enhanced the framework by developing a standardized pipeline for feature selection and normalization, aiming to further stabilize its performances across various datasets.

Our models showcased notably advanced performance compared to benchmark methods such as Scissor, scAB, and PACSI on two prostate cancer ST datasets. Notably, on the Erickson patient 1 dataset, our DEGAS predicted survival hazard scores demonstrated significant power in identifying tumor and TME regions com- pared to benchmark methods. Furthermore, on the 10x Genomics dataset, our DEGAS hazard scores exhibited higher consistency with cancer stage compared to benchmark methods.

Furthermore, we confirmed the DEGAS v2 model’s capability to discern disease- related differentiation of SCs within each cell type. In the Grubman and Mathys AD SC datasets, DEGAS scores revealed distinct patterns between SCs from AD patients and those from healthy individuals. This observation aligns with our hypothesis that cells from AD patients may undergo functional changes during disease progression and exhibit stronger disease associations compared to other cells of the same type. This method surpasses the limitations of approaches that focus only on alterations in cell type abundance or link disease to individual cell types.

Additionally, correlating disease scores predicted by various disease indices further underscored our model’s robustness in disease label selection.

Finally, we applied the DEGAS v2 model to the IUSM MM SC dataset, our model inferred that SC groups related to malignant disease outcomes exhibiting higher initial drug response, which was counter intuitive. However, all of these SC groups were associated with poor long-term drug responses, which revealed the underlying reason of their high hazard scores. This discovery highlighted the importance of considering longitudinal drug response in understanding clinical outcomes and emphasizing the need for novel drug discovery efforts.

We underscore that the robustness of our approach derives not solely from TL DL frameworks but also from our meticulous design of the feature selection and pre- processing pipeline.

However, feature selection remains a challenge, particularly due to the enormous feature dimension as well as their correlation. Both of these factors impede the effectiveness of Lasso or Elastic based-models. Therefore, integrating prior biological knowledge and advanced ML feature selection methods is imperative for constructing a more informative training dataset.

Furthermore, the incorporation of advanced survival loss functions into DL models offers a promising avenue to stabilize model performance.

